# Lgr5 Controls Extracellular Matrix Production By Stem Cells In The Developing Intestine

**DOI:** 10.1101/757641

**Authors:** Valeria Fernandez Vallone, Morgane Leprovots, Romain Gerbier, Didac Ribatallada-Soriano, Anne Lefort, Frédérick Libert, Gilbert Vassart, Marie-Isabelle Garcia

## Abstract

The Lgr5 receptor is a marker of intestinal stem cells (ISCs) that regulates Wnt/b-catenin signaling. In this study, phenotype analysis of knockin/knockout Lgr5-eGFP-IRES-Cre and Lgr5-DTReGFP embryos revealed that Lgr5 deficiency during Wnt-mediated cytodifferentiation results in amplification of ISCs and early differentiation into Paneth cells, which can be counteracted by in utero treatment with the Wnt inhibitor LGK974. Conditional ablation of Lgr5 postnatally, but not in adults, altered stem cell fate towards the Paneth lineage. Together, these in vivo studies suggest that Lgr5 is part of a feedback loop to adjust the Wnt tone in ISCs. Moreover, transcriptome analyses revealed that fetal ISCs generate their own extracellular matrix components, a property lost in adult ISCs, which adopt a definitive epithelialized phenotype and an inflammatory response signature. Absence of Lgr5 in fetal ISCs resulted in reduced extracellular matrix production and accelerated ISC maturation, indicating that Lgr5 regulates the ISC niche. Finally, evidences are provided that Rspondin 2 negatively regulates the pool of ISCs in organoids via Lgr5, revealing a sophisticated regulatory process for Wnt signaling in ISC.

## INTRODUCTION

The adult intestinal epithelium is a specialized tissue involved in nutrients absorption and protection against pathogens or environmental toxic agents. Under homeostatic conditions, within few days, this epithelium undergoes rapid and constant renewal supported by a pool of intestinal stem cells (ISCs), also called Crypt Base Columnar cells, identified by the expression of the Lgr5 receptor [1]. Restricted to the bottom of the crypts of Lieberkühn, ISCs have the capacity to both self-renew and give rise to transit amplifying cells, which differentiate along the villus architecture into all the cell lineages of the epithelium, i. e. absorptive Enterocytes, mucus-producing Goblet cells, hormones-secreting Enteroendocrine cells, Paneth cells generating antimicrobial products and the Type 2 immune response-inducer Tuft cells [2]. Other populations of slowly-cycling or label-retaining reserve stem cells have been identified able to efficiently regenerate the intestinal epithelium upon loss of Lgr5-expressing stem cells; additional evidences have been provided for coexistence and possible mutual interconversion between these two stem cell populations [3-5]. How the definitive crypt-villus architecture is reached in the adult intestinal epithelium and how adult stem cells emerge and establish from the embryonic gut tube deriving from endoderm, these are fundamental questions currently subject of active investigation. By using transgenic mouse lines, evidences have been provided that Cdx2 is a master transcription factor required for intestinal specification before the embryonic stage E14 [6-7]. Thereafter, the intestinal epithelium undergoes a profound remodeling, in part instructed by the underlying mesenchyme, leading to appearance of separate domains constituted by villus and intermingled intervillus regions [8-10]. Coherent with a proximal-to-distal wave of cytodifferentiation along the intestine mediated by the Wnt/b-catenin pathway around E14.5, the Wnt/β-catenin target gene Lgr5 becomes upregulated and identifies cells (ISC precursors) restricted to the intervillus regions that grow as adult-type organoids in the *ex vivo* culture system [11-14]. After birth, concomitant with Paneth cell lineage differentiation, intestinal crypts will be formed by invagination of the intervillus regions into the surrounding mesenchyme, bearing in their bottom, the Lgr5-expressing adult ISC [15].

Despite general consensus on the function of the Lgr5 receptor as a Wnt/β-catenin signaling modulator in stem cells, how it does so remains still controversial. First of all, *in vitro*, binding of the natural ligands Rspondins to the receptor Lgr5 has been demonstrated to either enhance or inhibit the Wnt pathway depending on the cell type analyzed [16-20]. Secondly, *in vivo*, homozygous *Lgr5-LacZNeo* knockin/knockout embryos deficient for Lgr5, exhibited an overactivated Wnt/b-catenin signaling pathway at birth associated with precocious Paneth cell differentiation, this suggesting a negative regulatory function of Lgr5 on this cascade [21]. However, conditional ablation of the Lgr5 function in adults did not result in significant alteration in Paneth cell differentiation [17]. Moreover, the molecular mechanisms associated with Lgr5 function in ISCs are still debated, does this G-protein-coupled receptor simply control Wnt signaling at the extracellular level by trapping the E3 ubiquitin ligase Znrf3/Rnf43 at the cell membrane or does Lgr5 signal *via* its transmembrane domains and intracellular tail [17,22-23] ?.

In the present report, we further investigated the role of the Lgr5 receptor during intestinal development by analyzing the transcriptome of Lgr5-expressing or Lgr5-deficient ISC just after the onset of the Wnt-mediated cytodifferentiation (E16) and in adult homeostatic tissues. We provide evidences that Lgr5 controls ISC maturation associated with acquisition of a definitive stable epithelial phenotype, that depends on the capacity of ISCs to generate their own extracellular matrix. In addition, using the *ex vivo* culture system, we demonstrate that the Rspondin 2 ligand negatively regulates the pool of ISCs in organoids via the Lgr5 receptor.

## RESULTS

### *In utero* inhibition of Wnt activity counteracts premature Paneth cell differentiation induced by Lgr5 deficiency in the intestine

To clarify the molecular function of the Lgr5 ISC stem cell marker in the embryonic intestine, we investigated the potential phenotype of knock-in/knockout (KO) homozygous Lgr5 embryos from the Lgr5-GFP-Cre^ERT2^ and Lgr5-DTReGFP mouse strains [1,24]. Since Lgr5 KOs generated from both transgenic lines showed neonatal lethality associated with ankyloglossia, histological analyses were performed at E18.5. Despite no evidence of gross architectural epithelial alterations, Lgr5 KOs exhibited early differentiation towards the Paneth lineage as revealed by Lendrum’s staining as well as qRT-PCR analysis of E18.5 tissues (Fig 1A and B, Fig S1), confirming previous studies on other Lgr5-deficient mouse strains [21,25]. In addition, quantification of the stem cell pool with the Olfm4 marker indicated expansion of this population in the intervillus (IV) region of Lgr5 KOs as compared to that of wild-types (WTs) (Fig 1C). Accordingly, Lgr5 KOs showed 4-fold increased expression of Wnt/β-catenin target genes (*Ascl2, Axin2*), histologically detected in the IV region (Fig 1D and E). Of relevance, upregulation of the truncated *Lgr5* transcript itself was even higher [10-fold versus (vs) WTs], suggesting a negative control of the Lgr5 receptor on its own expression (Fig 1D). Altogether, these data suggested that Lgr5 deficiency generates overactivation of the Wnt/β-catenin pathway in the prenatal small intestine that induces an expansion of ISC precursors and leads to premature Paneth cell differentiation around birth.

**Figure 1.**
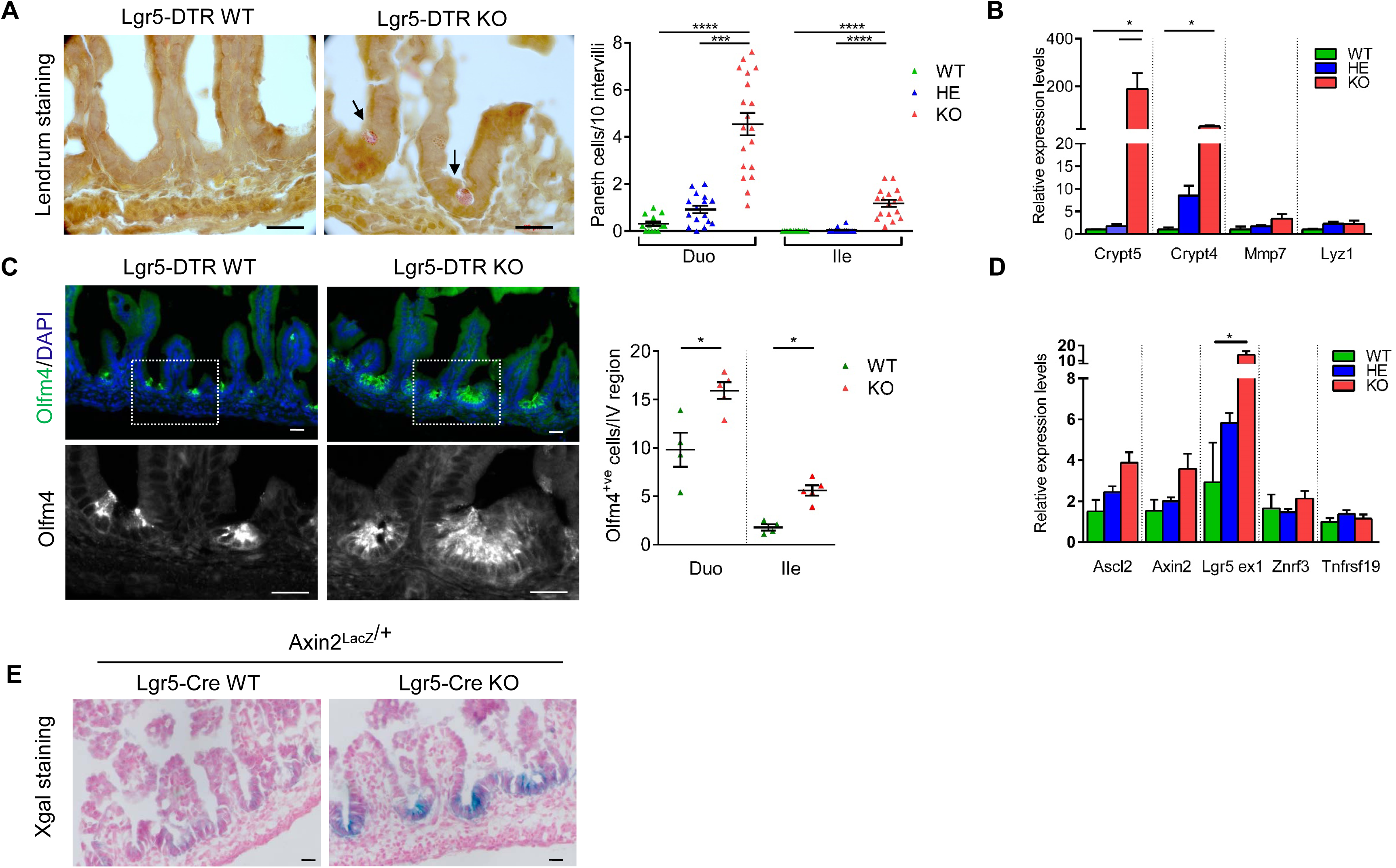
Lgr5 deficiency induces early Paneth cell differentiation and stem cell expansion in the small intestine at E18.5. A. Paneth cell quantification on Lgr5-DTReGFP duodenum (Duo) and Ileum (Ile). Left panel: Representative images of Lendrum’s staining. Arrows show differentiated Paneth cells. Right panel: quantification of the number of cells per 10 intervillus region. Each dot indicates the value for a given embryo. B. Expression analysis by qRT-PCR of the indicated Paneth cell markers in Lgr5-DTReGFP ileums (n= 3 WT, 6 HE, 6 KO). C. Immunofluorescence showing Olfm4^+ve^ cells in Lgr5-DTReGFP duodenums. Cell nuclei were counterstained with DAPI. Quantification of Olfm4^+ve^ cells per intervillus (IV) region in Duo and Ile. Each dot indicates the value for a given embryo D. Gene expression analysis by qRT-PCR of the indicated stem cell markers in Lgr5-DTReGFP ileums (n= 3 WT, 6 HE, 6 KO). E. Axin2/LacZ expression detected by X-gal staining in Axin2^Lac/+^-Lgr5-GFP-CreERT2 WT or KO ileums. Data information: All scale bars, 20 µm. Data are represented as means ± sem. *P< 0.05; ***P< 0.001; ****P< 0.0001 by Kruskal-Wallis test followed by Dunns multiple comparison test (A, B, D) and by Mann-Whitney test (C).

In attempts to reduce *in vivo* the excessive Wnt signaling tone observed in Lgr5 KO embryos, we treated pregnant females (Lgr5-DTReGFP and Lgr5-GFP-Cre^ERT2^ strains) with the orally administrable Wnt inhibitor LGK974. This inhibitor of the acyl transferase Porcupine (which alters Wnt ligand secretion) restores normal Wnt levels in tumor-bearing mice without affecting highly proliferative tissues such as the intestine [26]. Pilot experiments demonstrated that the compound efficiently crosses the placenta but that its administration before embryonic stage E11.5 can affect normal embryonic development (Fig S2A). Then, we tested two different administration windows for daily oral gavage (dose of 3 mg/kg/day), starting before (E13-E15) or during (E15-E17) the onset of Wnt-mediated cytodifferentiation (Fig 2A). When analyzed at E18.5, LGK974-treated tissues did not show any major alteration in the epithelium (Fig 2B). Administration of LGK974 between E15-E17, but not earlier, reduced Paneth cell differentiation in Lgr5 KOs to control levels (Fig 2C, Fig S2B). The treatment reduced expression of Wnt target genes (*Axin2, Ascl2, Lgr5 ex1*) expressed by ISCs without significantly altering expression of other reported stem cell markers (*Hopx, Tert*) (Fig 2D, Fig. S2C). LGK974 administration also induced downregulation of the Paneth cell marker *Crypt5* but not that of other cell lineages markers (Fig 2D, Fig. S2C). In contrast, attempts to upregulate Wnt/β-catenin signaling in *Lgr5*-expressing stem cells between E15-E17 by a genetic approach *via* deletion of the β-catenin exon 3 (encoding the sequences targeting the protein for proteasome degradation [27]) worsened the Paneth cell differentiation phenotype in Lgr5 KO embryos as compared to Lgr5 HEs (2.29 ± 0.33 vs 1.15 ± 0.22 paneth cells/10 intervilli, respectively; p=0.0317) (Fig S2D). Together, these rescue experiments further strengthened the notion that the Lgr5 receptor is involved in negative regulation of the Wnt/β-catenin activity at the onset of cytodifferentiation in the embryonic intestine.

**Figure 2.**
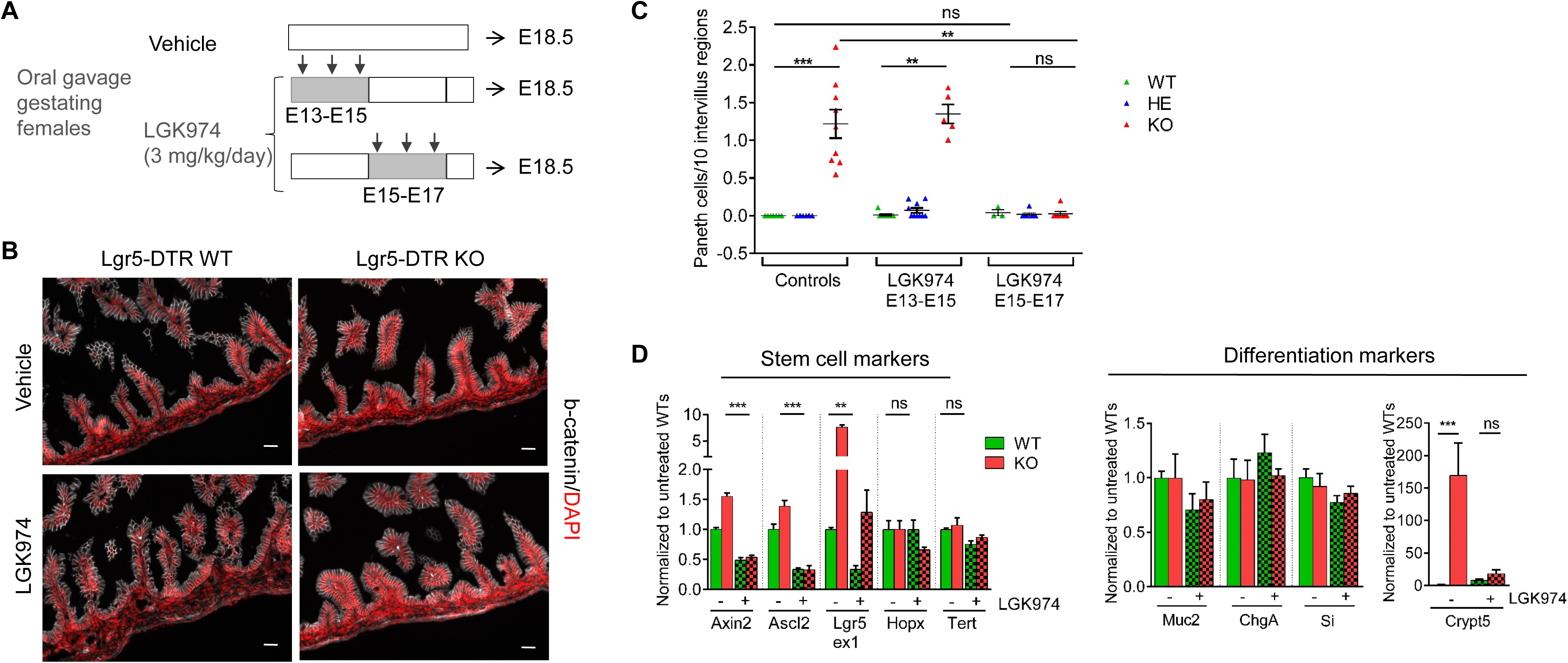
In utero inhibition of Wnt activity counteracts early Paneth cell differentiation induced by Lgr5 deficiency. A. Design of the experiment. Gestating females were vehicle- or LGK974-treated by oral gavage between E13-E15 or E15-E17. Small intestines of treated embryos were analyzed at E18.5. B. Immunofluorescence showing β-catenin expression in Lgr5-DTReGFP ileums. Cell nuclei were counterstained with DAPI. C. Paneth cell quantification on Lgr5-DTReGFP ileums. Each dot indicates the value for a given embryo (n= 3 to 11 embryos per condition). D. Gene expression analysis by qRT-PCR of the indicated stem cell and differentiation markers in Lgr5-DTReGFP ileums (vehicle-treated 7 WT and 7 KO; LGK974-treated 3 WT and 7 KO). Data information: All scale bars, 20 µm. Data are represented as means ± sem. ns not significant, **P< 0.01; ***P< 0.001 by Kruskal-Wallis test followed by Dunns multiple comparison test (C, D).

### Postnatal induction of Lgr5 ablation in ISCs alters stem cell fate towards the Paneth cell lineage

To determine whether the phenotype observed in Lgr5-deficient embryos could be reproduced postnatally (PN) when the Paneth cells normally emerge in control tissues, we generated conditional deficient-Lgr5 mice (cKO). These mice are double heterozygous Lgr5^GFP-CreERT2/flox^ in which cre-mediated deletion of Lgr5 exon 16 causes a frameshift and a null phenotype [1,17]. Following 3 consecutive tamoxifen injections to lactating females (PN days 6-8), we compared the fate of cKO Tom-recombined cells to that of control heterozygous (HE) Lgr5^GFP-CreERT2/+^/Rosa26R-Tom littermates after 10 days of chase (Fig 3A). No significant differences in terms of clone number were observed in Lgr5-ablated Tom^+ve^ tissues as compared to controls, suggesting that stemness was preserved during this period of chase (Fig 3B, C). However, cKO PN18 Lgr5-ablated Tom^+ve^ tissues exhibited a clear bias towards Paneth cell differentiation (Fig 3B and D). Such phenotype was observed in proximal and distal small intestines (Fig S3). When Lgr5 ablation was induced in adults already bearing a definitive number of Paneth cells in crypts, such phenotype was not observed 1 month after recombination; and no significant differences were detected in the proportion of traced clones (Fig 3B, D and E). Together, these data are consistent with the fact that loss of Lgr5 function in homeostatic adult tissues is not associated with an overt phenotype [17] meanwhile absence of the Lgr5 receptor during the prenatal and early postnatal stages impacts on stem cell fate.

**Figure 3.**
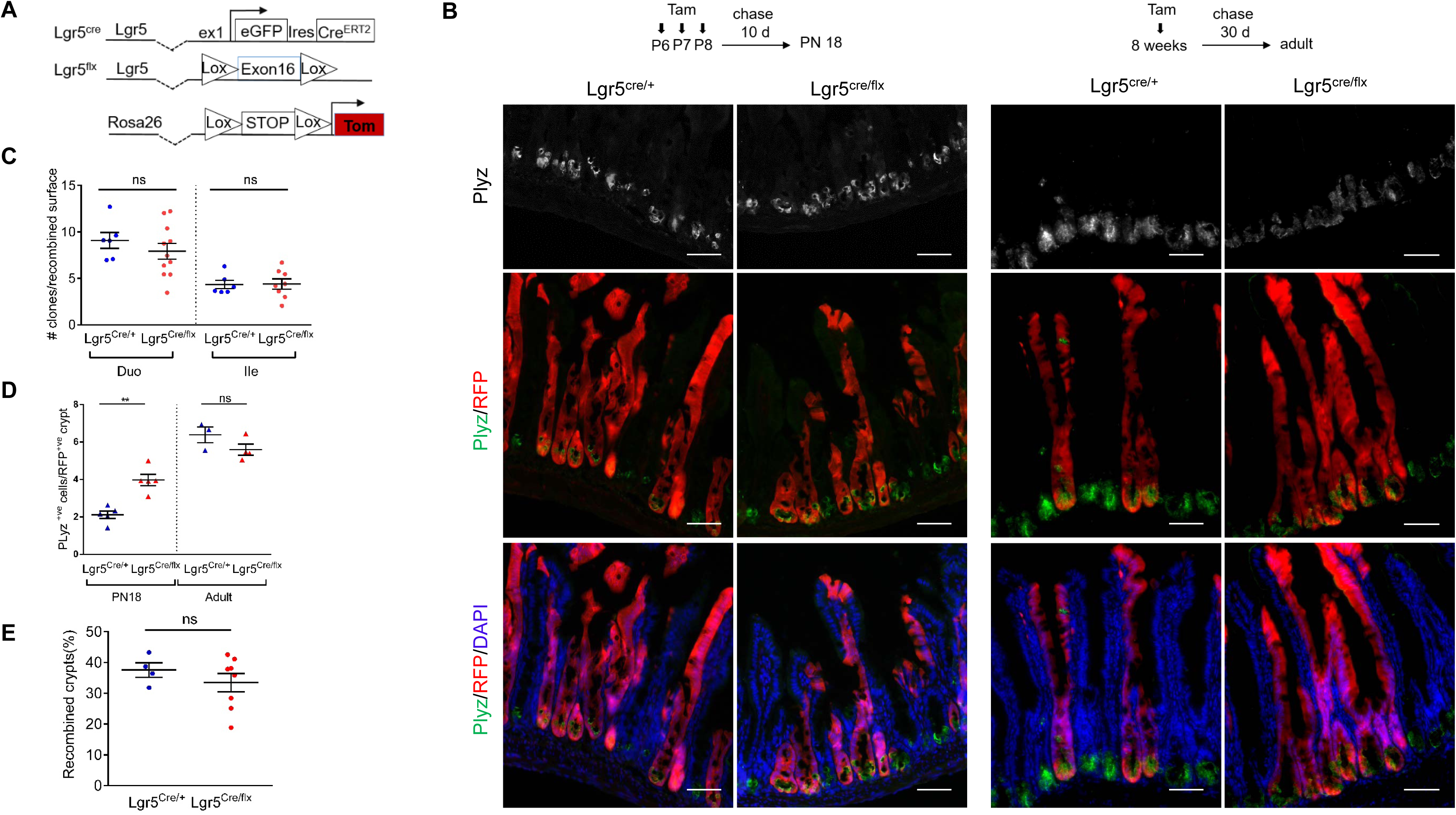
Postnatal Lgr5 ablation in ISCs alters stem cell fate towards the Paneth cell lineage. A. Schematic representation of the genetic elements for lineage tracing during postnatal development and adult homeostasis. B. Representative immunofluorescence pictures showing Paneth cells (Plyz^+ve^) in traced clones (RFP^+ve^) from control (Lgr5^Cre/+^) and cKO (Lgr5^Cre/flox^) ileums. Cell nuclei were counterstained with DAPI. Left panels: Lactating females were tamoxifen (Tam)-treated between PN 6 and PN 8. Small intestines of treated pups were analyzed 10 days after the last injection (PN day 18); Right panels: Adult mice received one single Tam intraperitoneal injection and ileums were analyzed 30 days later. C. Quantification of the number of traced clones per recombined surface in PN18-old mice. Each dot indicates the value for a given mouse. D. Quantification of the number of Paneth cell number per recombined RFP^+ve^ crypt on Control and cKO ileums in PN18 and adult mice. Each dot indicates the value for a given mouse. E. Quantification of the percentage of recombined crypts in adult mice. Each dot indicates the value for a given mouse. Data information: All scale bars, 20 µm. Data are represented as means ± sem. ns, not significant, **P< 0.01 versus controls by Kruskal-Wallis test followed by Dunns multiple comparison test (C). by Mann-Whitney test (D) and unpaired t test (E).

### Lgr5 controls extracellular matrix autocrine production in stem cells

To investigate the molecular pathways altered by Lgr5 deficiency, we analyzed the transcriptomic profile of GFP^+ve^ ISC precursors from Lgr5-DTReGFP HEs and KOs at E16.5. At this developmental stage, potential impact of Paneth cells was excluded based on Lendrum’stainings. Two pools of Lgr5 HEs (n= 7 and 8) and Lgr5 KOs (n= 2 and 6) coming from 2 different litters were isolated by Fluorescence-Activated cell sorting (FACS). RNAseq on these samples followed by differential gene expression analysis identified 487 genes (96 upregulated/391 downregulated) modulated in Lgr5 KOs vs HEs (False Discovery rate 0.1 and fold change above 1.5) (Fig 4A and B). Lgr5 deficiency was strikingly associated with de-enrichment in the Epithelial Mesenchymal Transition Gene dataset (47 genes modulated out of 200 genes in the EMT dataset, p value 4.e^-60^) (Fig 4B and C). Gene Ontology-Panther analysis of this short gene list indicated profound reorganization of the matrisome with significant reduction in Extracellular matrix (ECM) structural constituents, including collagen fibrils, in Lgr5-deficient ISC precursors (Fig 4B). In accordance with ECM playing a role in development [28], downregulated genes were also associated with tissue development, morphogenesis and regulation of cell migration (Fig 4B). Reduced expression of the Cadherin member Cdh11 and increased expression of the apical junction component Claudin 4 (Cldn4) suggested enforcement of an epithelial phenotype in Lgr5 KO precursors, coherent with evolution of these precursor ISCs towards a more mature stage (Fig 4A). To test this hypothesis, we compared the transcriptomes of E18.5 embryos and adult Lgr5 HE ISCs. Stem cell maturation was associated with downregulation of EMT processes with decreased expression of ECM-associated genes (Fig 4B, 4C and S4A). Moreover, in agreement with the study of Navis et al [29] investigating transcriptomic changes during suckling-to-weaning transition, maturation of ISCs also involved metabolic changes: with downregulation of the neonatal-related genes *Lactase (Lct), Arginosuccinate synthetase 1* (*Ass1*) and upregulation of the adult-related *Arginase 2* (*Arg2*) and *Sucrase-isomaltase* genes (*Si*) (Fig S4A). Together, these findings revealed that the transcriptome of ISCs substantially evolves during development, changing from an immature mesenchymal-like to a definitive mature epithelial phenotype. In addition, embryos deficient for Lgr5 demonstrate an accelerated conversion to an adult-type stage.

**Figure 4.**
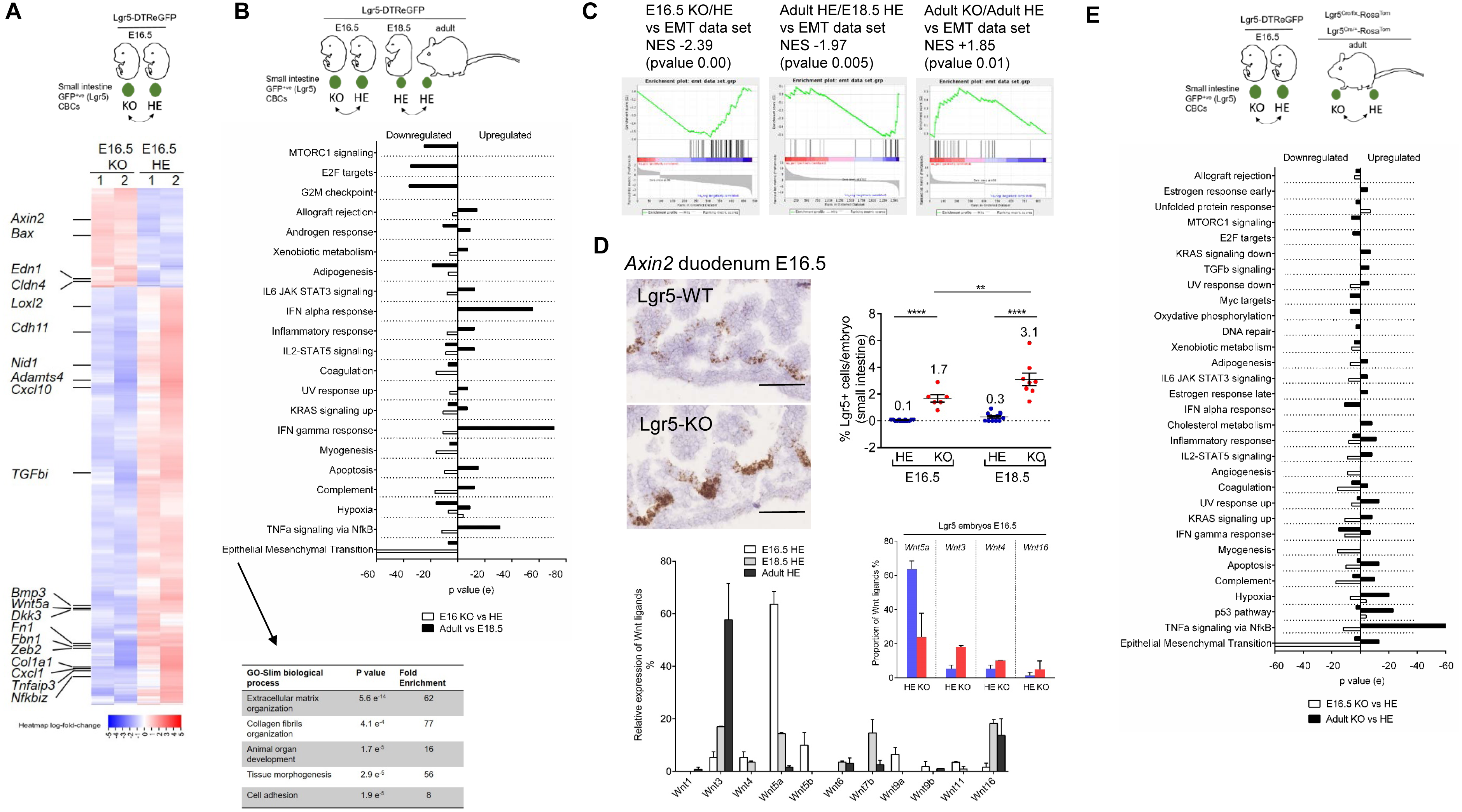
Transcriptome analysis of Lgr5-deficient ISC precursors. A. Design of the experiment. ISC (eGFP^+ve^) cells were sorted by FACS from Lgr5-DTReGFP embryos at E16.5 and RNA extracted was subject to RNAseq analysis. Heatmap of differentially regulated genes in the two independent pools (1,2) of HE and KO ISCs at E16.5. B. Design of the experiment. Compared Hallmark MSigDB gene set analysis on upregulated and downregulated genes in eGFP^+ve^ cells isolated from Lgr5 KOs vs Lgr5 HEs at E16.5 and Lgr5 adults vs E18.5 HEs. Below: GeneOntology analysis of downregulated genes identified in the Epithelial Mesenchymal Transition Gene Set. C. GSEA showing enrichment of the EMT data set in E16.5 Lgr5 KOs vs E16.5 HEs, adult HEs vs E18.5 HE embryos, and adult Lgr5 cKO vs Lgr5 HE modulated genes. NES: Normalized Enrichment Score. D. Upper left: in situ hybridization showing Axin2 expression in the duodenum of E16.5 Lgr5-DTReGFP embryos; Upper right: quantification of the number of eGFP^+ve^ (ISC) cells per small intestine at the indicated developmental stages in Lgr5 HEs and Lgr5 KOs. Each dot indicates the value for a given embryo. Below left: graph showing relative expression of the Wnt ligands at different developmental stages E16.5, E18.5 and adult HEs; Below right: graph showing relative expression of the main Wnt ligands in Lgr5 KOs and HEs at E16.5. E. Compared Hallmarks for upregulated and downregulated genes in the transcriptome of E16.5 Lgr5 KOs vs E16.5 HEs and Adult Lgr5-Cre cKO vs HEs. Data information: Scale bar, 100 µm. Data are represented as means ± sem. **P< 0.01; ****P<0.001 Two-way ANOVA followed by Tukey’s multiple comparisons test (D).

Transition from a mesenchymal to an epithelial phenotype is controlled by various signaling pathways [30]. Consistent with an involvement of the Wnt/β-catenin pathway, Lgr5 E16.5 KO GFP^+ve^ ISCs exhibited upregulation of the *Axin2, Bax* and *Edn1* target genes and the ISC number at E16.5 appeared significantly expanded in KOs as compared to HEs (Fig 4A and D, upper right panel). Relative levels of the most expressed Wnt Frizzled (Fzd5 and Fzd7) and Lrp coreceptors (Lrp6) were not significantly changed by the developmental stage of ISCs or the Lgr5 expression level (HE or KO) but expression of Wnt ligands varied over time (Fig 4D lower panels, Fig S4B). Indeed, expression of the non-canonical Wnt5a ligand, predominant in HE ISCs at E16.5 (in agreement with data from [13]); progressively dropped at a later prenatal stage (E18.5) and adulthood meanwhile expression of the canonical Wnt3 ligand increased in ISCs during the same period (Fig 4D, Fig S4A). In Lgr5-deficient ISCs, the Wnt5a-to-Wnt3 shift expression was detected at an earlier stage (E16.5) as compared to Lgr5 HEs pairs (Fig 4D, lower right panel). In addition, the TGFβ pathway involved in EMT was found deregulated in absence of the Lgr5 receptor with downregulation of *Bmp3* and *TGFβi* in E16.5 KOs as compared to HEs (Fig 4A). Moreover, whereas the inflammatory response (involving TNFα/NFkB and IFN α/γ pathways) was found upregulated during normal ISC maturation, this did not occur in E16.5 ISC KOs; these cascades appearing downregulated as compared to E16.5 ISC HEs (with downregulation of *Cxcl1, Tnfaip3, Cxcl10* and *Nfkbiz* genes) (Fig. 4A and B). Surprisingly, when Lgr5 function was specifically disrupted in adults using Lgr5^Cre/flox^ mice, most of the pathways downregulated in E16.5 KOs were instead upregulated (Fig 4E). As an exception, the IFN α/γ pathway appeared similarly downregulated in Lgr5-deficient embryo or adult stem cells (Fig 4E). Altogether, these data indicated that the transcriptome of ISC significantly changes from prenatal to adult stage, acquiring a definitive epithelial phenotype associated with: i) a net decrease in the capacity to generate its own extracellular matrix niche, ii) a conversion in Wnt ligand autocrine production and iii) an increased ability to respond to inflammatory signals. Moreover, absence of Lgr5 expression in the early ISC precursors (i.e. just after their emergence in intervillus regions) leads to accelerated conversion of ISCs towards their epithelial mature phenotype whereas acute ablation of Lgr5 in adults rather shifts epithelial stem cells towards a mesenchymal-like phenotype.

### Stemness is preserved in Lgr5-deficient organoids *ex vivo*

ISC co-express the two paralogue receptors Lgr4 and Lgr5 [17,31]. Since deficiency for the Lgr4 receptor leads to ISC loss due to insufficient Wnt signaling in cultured crypts, we assessed the long-term growth properties of Lgr5-deficient ISCs in the *ex vivo* culture system [31]. Irrespective of the mouse strain of origin, Lgr5 KO E18.5 small intestines generated a 3-4-fold increase in the absolute number of growing organoids which exhibited higher complexity (measured by the branching coefficient) as compared to WTs and HEs upon initial seeding (Fig 5A). Then, we assessed whether the Wnt tone in Lgr5 KO organoids would be sufficient to confer reduced growth requirements for the Rspondin 1 (Rspo1) ligand. Despite an overall 2-fold increase in Wnt signaling activity (*Axin2, Ascl2, Olfm4, Lgr5*) at any Rspo1 concentration tested as compared to Lgr5-DTReGFP WTs, KO organoids remained dependent on Rspo1 to grow *ex vivo* (Fig 5B and C). As observed in tissues, differentiation towards the Paneth cell lineage was increased in Lgr5 KO cultures (Fig S5A). Since transcriptomic analysis of Lgr5 KOs had suggested modifications in the stemness status of precursor ISCs, we studied whether this might affect long-term maintenance of ISCs *ex vivo* by passaging Lgr5-DTReGFP samples for more than 20 passages (Fig S5B). Expression of Wnt target genes was maintained over passages as compared to Lgr5 WTs (Fig S5C). Together, these results indicated that long-term replating of Lgr5 KO organoids is preserved *ex vivo*, without evidence of Lgr5 deficiency leading to loss of stemness.

**Figure 5.**
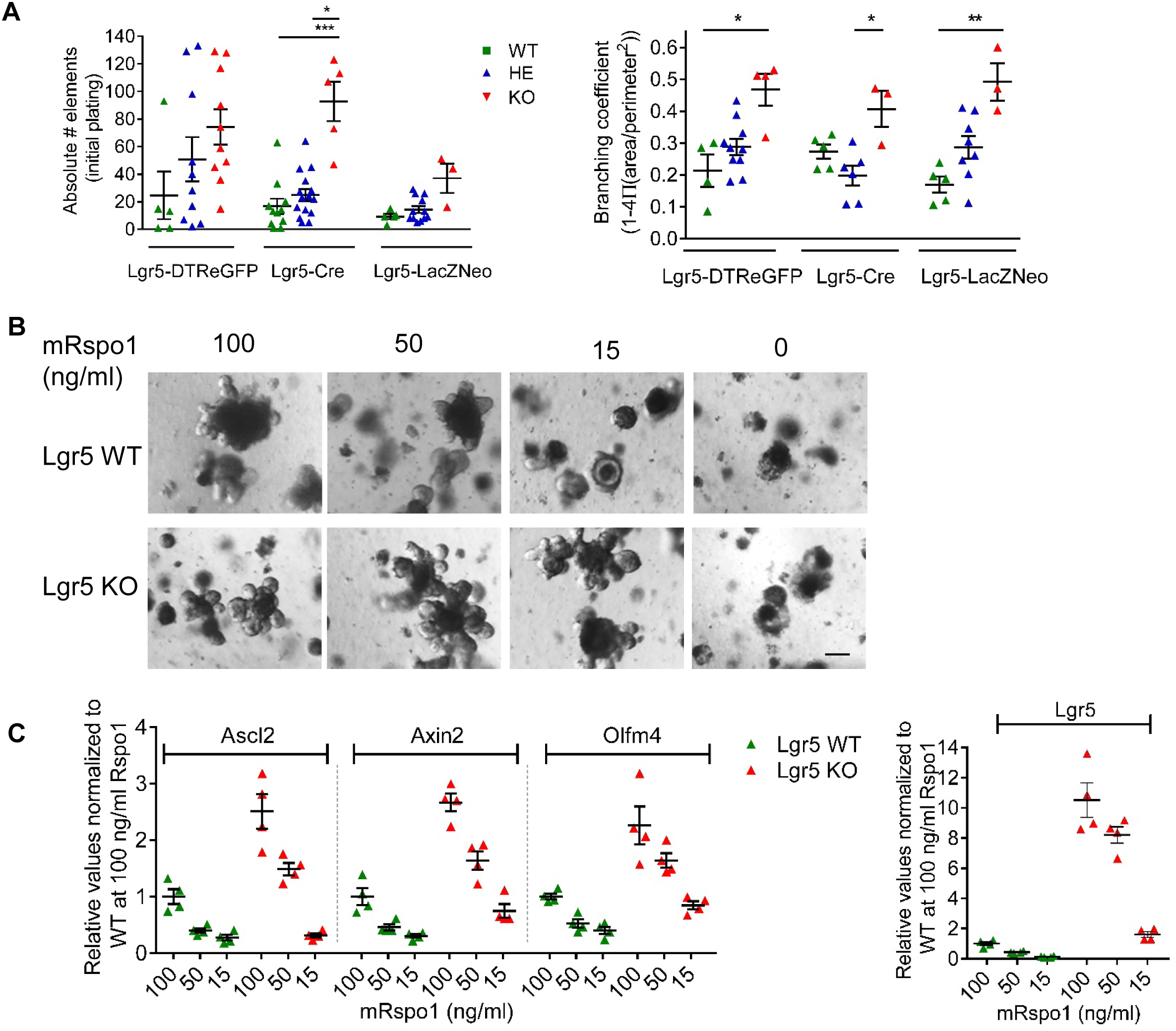
Stemness is preserved in Lgr5-deficient organoids. A. Plating efficiency and organoid complexity of ex vivo cultured E18.5 small intestines (the number of embryos analyzed is indicated in the graph). Pictures were acquired 6 days after initial seeding. B. Influence of mouse Rspondin 1 concentration on growth of replated Lgr5-DTReGFP WT and Lgr5 KO organoids after 5 days of culture. C. Gene expression analysis by qRT-PCR of the indicated stem cell markers in Lgr5-DTReGFP WT and Lgr5 KO organoids at the indicated Rspondin 1 concentrations after 5 days of culture. Each dot indicates the value for an organoid culture originating from a given embryo. Data information: Scale bar, 50 µm. Data are represented as means ± sem. *P<0.05; **P< 0.01; ***P<0.001. Kruskal-Wallis test, followed by Dunn’s multiple comparison test (A, C).

### Rspondin 2 ligand/ Lgr5 receptor interaction regulates stem cell fate in organoids *ex vivo*

Rspondins (Rspo1/Rspo2/Rspo3/Rspo4) have been reported to behave as redundant ligands for Lgr receptors, leading to enhanced Wnt/β-catenin activity [16-20]. We investigated the impact of Rspondin members on organoid growth *ex vivo* by culturing Lgr5-DTReGFP WTs isolated from E18.5 small intestines in the Sato medium containing EGF, Noggin plus either mRspo1 or mRspo2. Both conditions allowed efficient growth of Lgr5 WT organoids after 5 days of culture (Fig 6A). However, when cultured in Rspo2 medium, organoids emitted elongated protrusions as compared to those grown in the Rspo1 medium (Fig 6A). As assessed by qRT-PCR analysis, culture of WT organoids in mRspo2 conditions induced downregulation of the stem cell markers *Ascl2* (by 70%), *Axin2* and *Olfm4* (by 30%) as well as the *Lgr5* gene itself (by 40%) as compared to mRspo1 conditions (Fig 6B). Accordingly, *in situ* hybridization and immunohistochemistry experiments confirmed decreased *Axin2* expression in crypt-like domains and reduction in the proportion of Olfm4-expressing cells in WT organoids in Rspo2 vs Rspo1 conditions (Fig 6C). In contrast to WT organoids, KO organoids did not demonstrate such morphological differences in Rspo1 and Rspo2-containing media (Fig 6A). Wnt target gene expression remained at higher levels in Lgr5 KO organoids as compared to WTs irrespective of the Rspondin type used for culture (Fig 6B and C). Therefore, these data indicated that Rspo ligands do not exhibit redundant functions on ISCs grown *ex vivo*, this likely involving the Rspo2/Lgr5 axis. Further, to determine whether this interaction in ISCs can also affect cell differentiation, we analyzed the expression of specific cell-lineage markers by qRT-PCR (Fig 6D). Replacement of Rspo1 by Rspo2 did not impact on ChgA^+ve^ enteroendocrine or Muc2^+ve^ Goblet cell differentiation, but induced a bias towards the SI^+ve^ absorptive lineage to the expense of the Crypt5^+ve^ Paneth cell type in WT organoids (Fig 6D). Analysis of Lgr5-deficient organoids suggested that the Rspo2/Lgr5 interaction might control commitment towards the absorptive fate but also that Rspo2 can regulate Paneth cell differentiation *via* its interaction with cell surface molecules other than the sole Lgr5 receptor.

**Figure 6.**
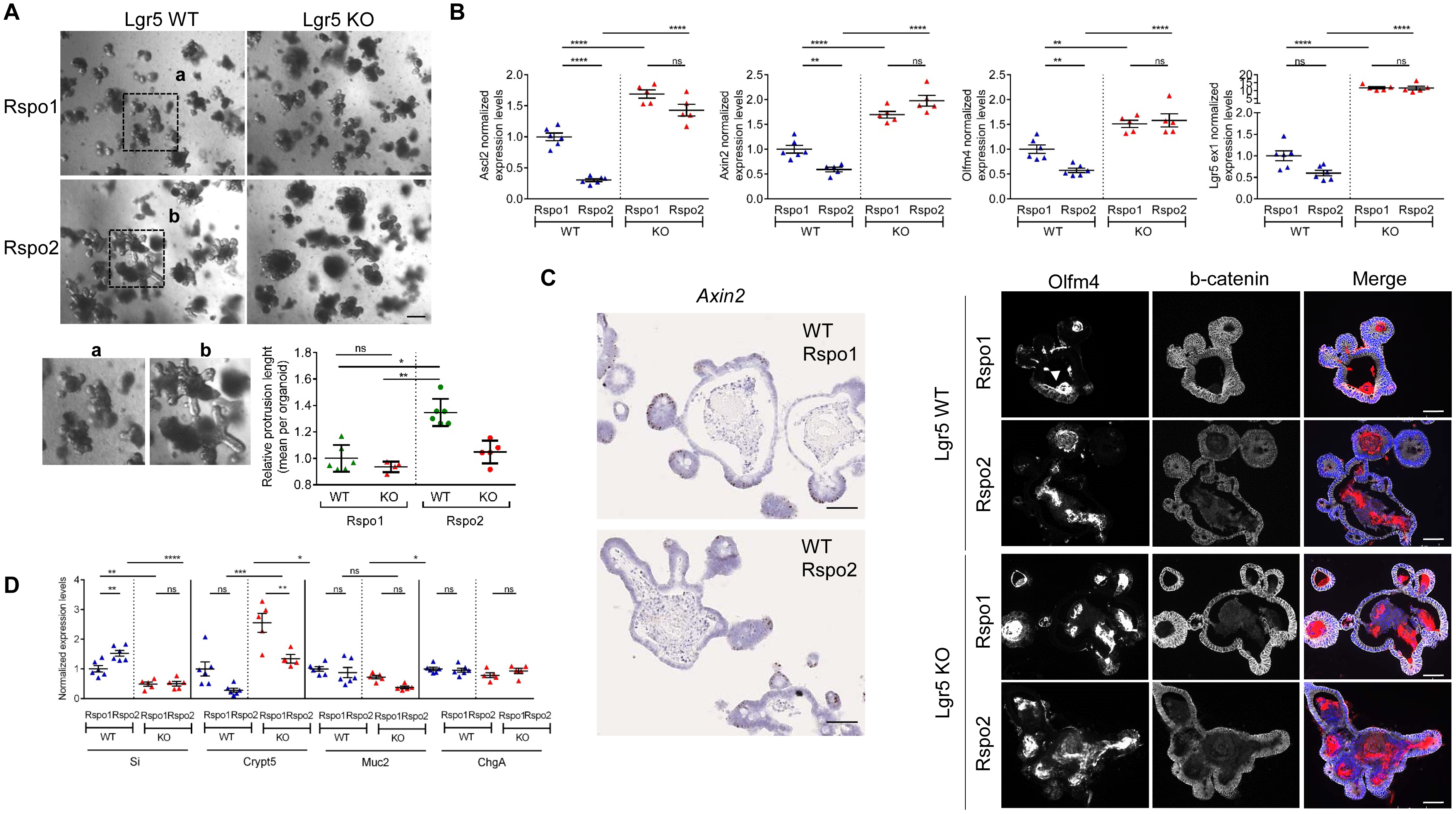
Rspondin 2/Lgr5 interaction regulates stem cell fate in organoids. A. Influence of Rspondin type (Rspo1 or Rspo2) on growth of replated Lgr5-DTReGFP WT and Lgr5 KO organoids. Insets a and b correspond to higher magnification of the representative pictures showing elongated crypt-like domains in Lgr5 WT/Rspo2 cultures. Quantification of the differences in morphology of organoids after 5 days of culture. B. Gene expression analysis by qRT-PCR of the indicated stem cell markers in Lgr5-DTReGFP WT and Lgr5 KO organoids after 5 days of culture. Each dot indicates the value for an organoid culture originating from a given embryo. C. Left panel: in situ hybridization showing Axin2 expression in Lgr5-DTReGFP WT and KO organoids cultured in Rspo1 or Rspo2-containing medium; right panel: immunofluorescence showing Olfm4^+ve^ cells in Lgr5-DTReGFP WT and KO organoids. Epithelium is delineated with β-catenin and nuclei counterstained with DAPI (Merge). The white arrowhead evidences accumulated Olfm4 in the luminal side of organoids. D. Gene expression analysis by qRT-PCR of the indicated lineage specific markers in Lgr5-DTREeGFP WT and KO organoids after 5 days of culture. Each dot indicates the value for an organoid culture originating from a given embryo. Data information: Scale bars, 100 µm (A) and 50 µm (C). Data are represented as means ± sem. ns= not significant; *P< 0.05; **P< 0.01; ***P< 0.001. by Kruskall-Wallis test followed by Dunns multiple comparison test (A); and by Two-way ANOVA followed by Tukey’s multiple comparisons test (B,D).

## DISCUSSION

Since the identification of the atypical GPCR receptor Lgr5 as a bone-fide stem cell marker in the normal intestinal epithelium as well as in cancer cells, the potential biological function of this receptor has been questioned but still, it remains elusive and controverted [32]. The biological function of Lgr5 was initially addressed in Lgr5-null embryos using the Lgr5-LacZNeo mouse line, in which exon 18 coding for the transmembrane and intracellular tail, is replaced by the reporter cassette [33]. The observed phenotype suggested that Lgr5 exerts a negative regulatory role on the Wnt/β-catenin pathway in stem cells [32]. Following the publication of reports identifying Rspondins as ligands for Lgr5, and showing that this interaction enhances Wnt/β-catenin signaling *in vitro*, we have readdressed the biological relevance of this marker on two other plain knock in/knockout Lgr5eGFP-Ires-CreERT2 and Lgr5-DTR mouse strains, which differ from the Lgr5-LacZNeo one by the fact that reporter cassettes are inserted within the first exon of Lgr5 [1,17]. Irrespective of the mouse strain considered, the observation that Wnt ligand inhibition at the onset of cytodifferentiation could interfere with the precocious Paneth cell differentiation observed at birth in knockouts, further strengthened conclusions drawn from the first report on a role of Lgr5 in negative regulation of the Wnt/β-catenin activity during the prenatal stage. Similarly, conditional ablation of the Lgr5 function in the postnatal phase during which Paneth cell differentiation normally proceeds, accelerated the maturation process. In adults, no overt phenotype was observed upon Lgr5 loss (though significant transcriptomic changes were detected), likely blurred in a context where Paneth cells are fully established. In addition, phenotypic differences observed in fetal and adult tissues upon Lgr5 loss may also be explained by co-expression of the paralogue Lgr4 receptor demonstrated to play a dominant role in ISCs on Wnt signaling over Lgr5 [17,25,31]. Considering that Lgr5, but not Lgr4, is a direct Wnt/β-catenin target gene, one hypothesis is that both receptors might play complementary roles in ISCs, Lgr4 and Lgr5 regulating Wnt signaling under normal homeostatic and Wnt overactivated situations, respectively.

To investigate the molecular mechanisms associated with premature Paneth cell differentiation in Lgr5-deficient embryos at birth, the transcriptomes of heterozygous (control) and homozygous (KO) ISCs were compared at E16.5. Deficiency for Lgr5 induced a profound downregulation of many extracellular matrix (ECM) components i.e. collagens, proteoglycans (versican, decorin,), glycoproteins (laminin a4, fibronectin, thrombospondins, nidogens) and ECM-associated modifying proteins (Adamts proteases, Lysyl oxidase, cytokines, glycosaminoglycan-modifying heparan sulfate sulfotransferase), all involved in matrisome formation and maintenance. The highly dynamic matrisome, defined by a core of ∼ 300 proteins localizing at the basement membrane (between epithelial cells and stromal cells) and in the neighboring interstitial space, is known to regulate tissue development, as well as fibrosis and cancer progression in adults [28]. Our findings provide evidences that early ISC precursors (E16.5) are themselves able to synthesize their own ECM components, in particular collagen fibrils known to control matrix stiffness, this conferring to these epithelial cells some kind of mesenchymal-like properties. Coherent with a recent report demonstrating that softening of the matrigel in the *ex vivo* culture system enhances stem cell differentiation in adult ISC-derived organoids, we observed that decreased expression of ECM proteins by E16.5 KO ISCs, likely reducing stiffness of the tissue surrounding ISCs, correlated with precocious differentiation of Paneth cells before birth [34]. Moreover, according to our transcriptomic data on adult vs embryonic ISCs, the capacity of stem cells to synthesize their own ECM components progressively decreases during maturation, correlating with acquisition of a definitive “fully” epithelialized phenotype. In the adult intestinal crypt, the ISC microenvironment substantially differs from the prenatal intervillus domains, in particular through its cellular niche composed of adjacent epithelial Paneth cells as well as various resident stromal cell types, which all might contribute to provide the adequate ECM environment to ISCs [14,35-36]. Nevertheless, based on our transcriptome analyses of ISCs depleted of Lgr5 (Lgr5^Cre/flx^ vs Lgr5^Cre/+^), adult ISCs still conserve the control on their epithelial status as loss of the Lgr5 receptor favored transition from an epithelial to a more mesenchymal-like phenotype (EMT Hallmark upregulated in KOs vs HEs). In line with these findings obtained under *in vivo* homeostasis, Lgr5 knockdown on cultured colorectal cancer cell lines also leads to upregulation of EMT-related genes *in vitro* [37]. These results, linking Lgr5 function to EMT control in adult ISCs, highlight the interest to conduct further studies addressing the role of this receptor on the metastatic potential of cancer cells. Our study also sheds new lights regarding molecular mechanisms associated with ISC maturation. Indeed, in addition to Wnt signaling, other pathways were identified to be modulated in ISCs during this process, in particular the inflammatory response that was induced in adults vs E18.5 ISCs. Remarkably, in E16.5 Lgr5-null ISCs, transcriptome data suggested that the TNFα, and IFNα/γ pathways, well known to crosstalk with the Wnt cascade in a complex way, appeared downregulated as compared to control ISCs [38-39]. Future studies will be needed to determine if ISC maturation in Lgr5-null embryos, is not only accelerated but also improperly executed in this regard.

Initial *in vitro* studies, identifying Rspondins (Rspo1/Rspo2/Rspo3/Rspo4) as ligands for the three Lgr paralogues, suggested both ligand and receptor redundancy [16-19]. However, evidences have been provided that Rspo2, but not other members of this family, exhibits tumor suppressive activity on colorectal cancer cell lines *via* negative regulation of the Wnt/β-catenin pathway through interaction with the Lgr5 receptor [20]. In order to clarify this point, we compared the effect of Rspo1 and Rspo2 on ISC growth and differentiation in the *ex vivo* culture system. As compared to Rspo1, Rspo2 downregulated expression of ISC gene markers (*Ascl2, Olfm4, Lgr5, Sox9, Axin2*) in WT organoids, and differentially modulated cell lineage commitment (essentially affecting the Absorptive and Paneth lineages). The observation that ISC markers remained at similar higher levels in Lgr5-deficient organoids irrespective of the Rspondin used as compared to WTs, this is consistent with Rspo2 ligand/Lgr5 receptor interaction specifically negatively controlling the Wnt/β-catenin pathway in ISCs. Nevertheless, the observed effect of Rspo2 vs Rspo1 towards the Paneth cell lineage on Lgr5-null organoids might also be explained by interaction of Rspo2 with other receptors, possibly the cognate Lgr4 receptor or likely heparan sulfate proteoglycans, such as Glypican or Syndecan receptors (namely Gpc3/Gpc4 and Sdc1/Sdc4 that are the most expressed in ISCs-Fig S4B). These latter cell surface molecules have recently been reported as alternative receptors for Rspo2 and Rspo3 to regulate Wnt/β-catenin signaling [40-41]. *In vivo*, the main source of these ligands are peri-cryptal myofibroblasts, which predominantly express Rspo3 under normal conditions and can overexpress Rspo2 upon infection or inflammatory stimuli [42-43]. Considering this complex ligand expression pattern and that of the known receptors, it is tempting to speculate that regulation of the ISC fate through Rspo/Lgrs interaction might depend in part on both spatial and temporal release of Rspo ligands in the ISC niche.

## MATERIAL AND METHODS

### Experimental animals

Animal procedures complied with the guidelines of the European Union and were approved by the local ethics committee under the accepted protocols 535N and 631N. Mice strains were: *Lgr5*-DTR knock-in [24], Ctnnb1^exon3^ [27], Lgr5-LacZNeo [33], Lgr5-Flox exon16 [17], Lgr5-GFP-Cre^ERT2^ [1]. Rosa26CAG floxed stop tdTomato referred as Rosa26R-Tomato and *Axin2*-LacZ (Jax mice). The day the vaginal plug was observed was considered as embryonic day 0.5 (E0.5).

For LGK974 rescue experiments, pregnant females were administered the LGK974 compound (kindly provided by Novartis) by oral gavage once a day at a dose of 1-3 mg/kg/day.

For lineage tracing experiments, tamoxifen (Sigma-Aldrich) was dissolved in a sunflower oil (Sigma-Aldrich)/ethanol mixture (9:1) at 10 mg/ml and used at a dose of 0.1 mg/g of body weight for gestating and lactating females or at a single dose of 2 mg for adult mice by intra-peritoneal injection.

### Tissue processing and immunohistochemical analysis

Small intestine samples were immediately fixed with 10% formalin solution, neutral buffered (Sigma-Aldrich) overnight at +4°C and then sedimented through 30% sucrose solution before OCT embedding. Histological and X-gal staining protocols as well as immuno-fluorescence/histochemistry experiments on 6 µm sections were carried out as previously described [21]. Lendrum’s staining was performed according to the manufacturer’s instructions (cat # 631340, Clinitech, UK). The primary antibodies used for staining were: rabbit anti-Olfm4 (Cell Signaling), rabbit anti-Plyz (Dako), rat anti-RFP (Chromotek), mouse anti-beta-catenin BD). Samples were visualized with Zeiss Axioplan 2 (immunohistochemistry) or Zeiss Observer Z1 microscope (immunofluorescence).

Quantification of the number of Paneth cells per 10 intervillus units (IV) was performed on a mean of 50 IV in E18.5 tissues, on a mean of 20 and 100 Tomato-recombined crypts in postnatal and adult samples. Postnatal and adult lineage tracing analysis was performed on a mean of 100 crypt-villus units. Quantification of Olfm4^+ve^ cells per IV was performed on a minimum of 20 IV per embryo. The number of animals used for each experiment is reported in Figure legends.

### Ex vivo culture

Embryonic small intestine was dissociated with 5 mM EDTA-in DPBS (Gilbco) according to the protocol reported in [44]. Briefly, the culture medium used consisted in Advanced-DMEM/F12 medium supplemented with 2 mM L-Glutamine, N2 and B27 w/o vit.A (Invitrogen), gentamycin, penicillin-streptomycin cocktail, 10 mM HEPES, and 1 mM N acetyl cysteine. Growth factors were added at a final concentration of : 50 ng/ml EGF and 100 ng/ml Noggin (both from Peprotech), and 100 ng/ml CHO-derived mouse R-spondin 1 or 5 ng/ml CHO-derived mouse R-spondin 2 (R&D System). The final concentration of Rspondins in the culture medium was initially tested in pilot experiments in order to add comparable amounts of bioactive ligands (bioactivity measured by TOPflash assays and organoid growth curves, R&D systems). Culture medium was changed each other day and after 5-6 days in culture, organoids were harvested, mechanically dissociated and replated in fresh Matrigel (BD Biosciences). Culture media were supplemented with 10-µM Y-27632 (Sigma Aldrich) in all initial seeding and replating experiments. Pictures were acquired with a Moticam Pro camera connected to Motic AE31 microscope.

### Gene expression analysis

qRT-PCR was performed on total RNA extracted from embryonic tissues or organoid cultures as reported [21]. Expression levels were normalized to that of reference genes (Rpl13, Gapdh, Ywhaz). Each sample was run in duplicate. Primer sequences are reported in Table S1. *In situ* hybridization experiments were performed using the MmAxin2 RNAscope probe (cat # 400331, ACD, UK) according to manufacturer instructions with the RNAscope kit.

### RNA seq and Gene Set Enrichment Analysis (GSEA)

RNA quality was checked using a Bioanalyzer 2100 (Agilent technologies). Indexed cDNA libraries were obtained using the Ovation Solo RNA-Seq System (NuGen) following manufacturer recommendation. The multiplexed libraries were loaded on a NovaSeq 6000 (Illumina) using a S2 flow cell and sequences were produced using a 200 Cycle Kit. Paired-end reads were mapped against the mouse reference genome GRCm38 using STAR software to generate read alignments for each sample. Annotations Mus_musculus.GRCm38.90.gtf were obtained from ftp.Ensembl.org. After transcripts assembling, gene level counts were obtained using HTSeq. Genes differentially expressed were identified with EdgeR method (FDR 0.1 and Fold change 1.5) and hallmarks were analyzed using GSEA MolSig (Broad Institute) [45]. Then, Gene pattern was used to compare by pre-ranked GSEA E16.5 Lgr5 KO vs HE; Adult HE vs E18.5 HE, or Adult Lgr5 KO vs HE transcriptomes with Gene data sets [46]. Gene Ontology was used to identify biological processes [47]. Transcript profiling: GEO accession number will be communicated upon publication.

### Statistical analysis

Statistical analyses were performed with Graph Pad Prism 5. All experimental data are expressed as mean ± s.e.m. The significance of differences between groups was determined by appropriate parametric or non-parametric tests as described in Figure legends.

## Supporting information

Table S1 list of qPCR primers

## Acknowledgments

We are grateful to Genentech for providing us with the *Lgr5*-DTReGFP mice, Novartis for providing LGK974 compound, Makoto Taketo for providing the Ctnnb1 exon3 strain and Hans Clevers for providing the Lgr5-LacZNeo and Lgr5 flox mouse strains.

Grant support: supported by the Interuniversity Attraction Poles Programme-Belgian State-Belgian Science Policy (6/14), the Fonds de la Recherche Scientifique Médicale of Belgium, the Walloon Region (program CIBLES) and the non-for-profit Association Recherche Biomédicale et Diagnostic.

## Author contributions

VFV, study concept and design, acquisition of data, analysis and interpretation of data, statistical analysis, drafting of the ms

ML, RG, DRS, AL, FL: acquisition of data, analysis and interpretation of data, statistical analysis

GV: study concept and design, critical revision of the ms, obtained funding, study supervision

MIG: study concept and design, acquisition of data, analysis and interpretation of data, drafting of the ms, critical revision of the ms, study supervision

