## Supplementary figures for "Lgr5 Controls Extracellular Matrix Production By Stem Cells In The Developing Intestine"

**SUPPLEMENTARY DATA**

**Figure S1. Lgr5 deficiency induces early Paneth cell differentiation and stem cell expansion in the small intestine at E18.5.**
A. Paneth cell quantification on Lgr5-Cre duodenum (Duo) and Ileum (Ile). Quantification of the number of cells per 10 intervillus (IV) regions. Each dot indicates the value for a given embryo (Duo: n= 3 WT, 4 HE, 4 KO; Ile: n= 3 WT, 5 HE, 4 KO). Data are represented as means ± sem. *P< 0.05; by Kruskal-Wallis test followed by Dunns multiple comparison test.

B. Sagittal sections of craniofacial region showing the presence of an ankyloglossia in Lgr5-null Cre embryos at E18.5 as compared to the WT littermate. The tongue (T) and mandible (M) are indicated.

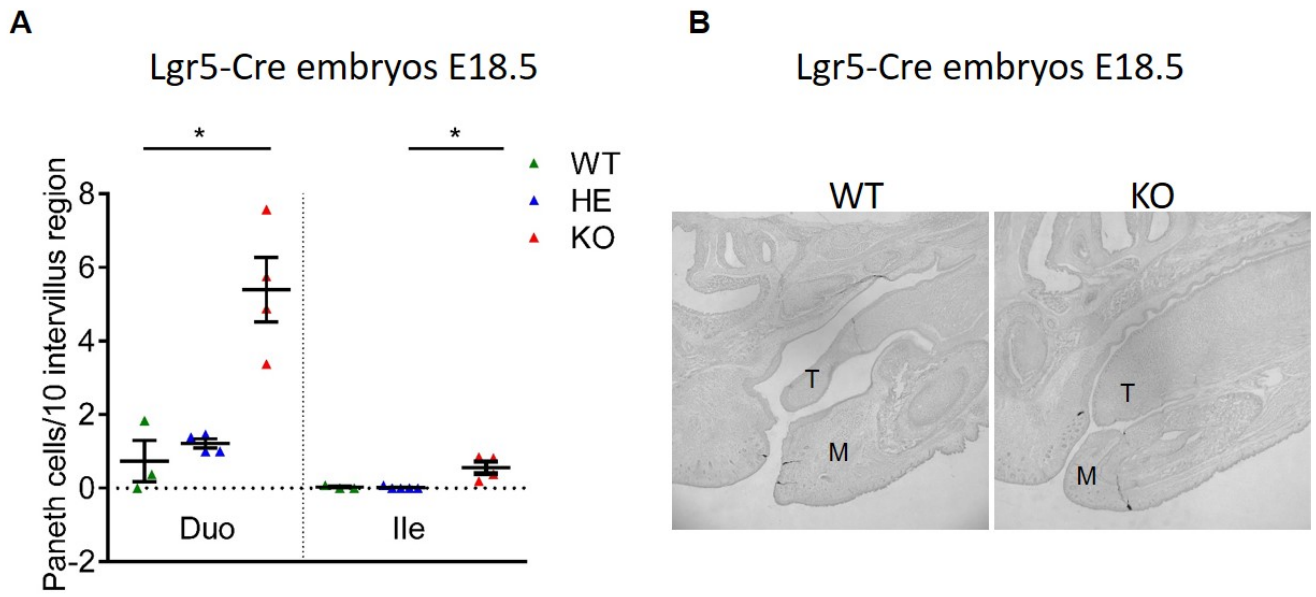

**Figure S2. In utero inhibition of Wnt activity counteracts early Paneth cell differentiation induced by Lgr5 deficiency.**

A. Gestating Lgr5-Cre heterozygous females were vehicle- or LGK974-treated by oral gavage between E9.5-E15 at the indicated dose. Global morphological analysis of whole embryos was done at E16.5.

B. Gestating Lgr5-DTReGFP heterozygous females were vehicle- or LGK974-treated by oral gavage between E13-E15 or E15-E17 at the indicated dose. Duodenum of treated Lgr5-DTReGFP null embryos were analyzed at E18.5 for Paneth cell differentiation. Each dot indicates the value for a given embryo.

C. Gestating Lgr5-Cre heterozygous females were vehicle- or LGK974-treated by oral gavage between E15-E17 at the indicated dose. Gene expression analysis of the indicated stem cell (*Axin2*) and Paneth differentiation (*Crypt5*) markers was performed at E18.5 in ileums of Lgr5-DTReGFP embryos by qRT-PCR. Each dot indicates the value for a given embryo.

D. Gestating females were intra-peritoneally injected with tamoxifen between E15-E17. Upper panel: ileums of embryos were analyzed at E18.5 for Paneth cell differentiation. Each dot indicates the value for a given embryo. Lower panel: Recombination of floxed b-catenin exon 3 was verified by PCR on ileums of embryos. Genotypes of embryos for b-catenin (Ctnnb1) and Lgr5 loci are indicated.

Data are represented as means ± sem. *P< 0.05; ***P< 0.001. by Kruskall-Wallis test followed by Dunns multiple comparison test (B-D).

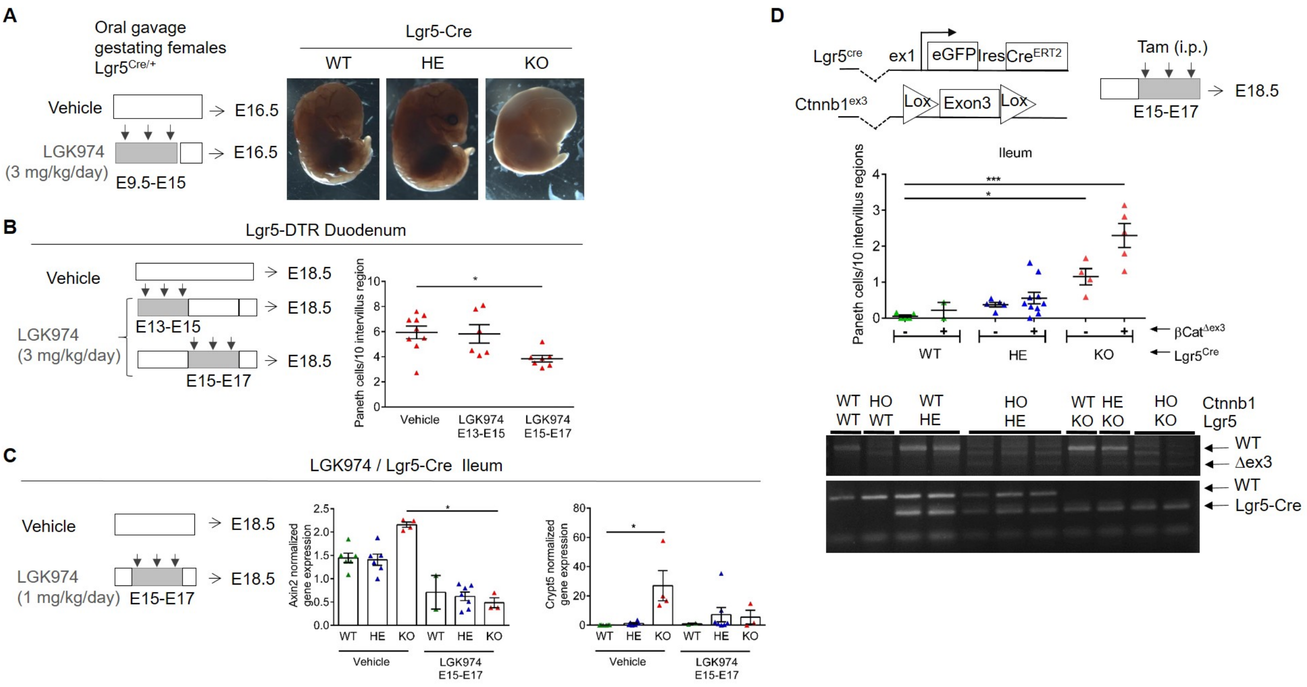

**Figure S3. Postnatal and adult Lgr5 ablation in ISCs alters stem cell fate towards the Paneth cell lineage.**

Fate of RFP$^{+ve}$-traced clones in control and cKO ileums at day PN18 after 10 days of chase. The proportion of fully recombined and mosaic RFP$^{+ve}$ crypt/villus units as well as RFP$^{+ve}$ cells only labelling Paneth cells is indicated for the duodenum and ileum of controls (Lgr5-Cre/+) and cKO (Lgr5-Cre/flx) mice (n= 5 for each genotype).

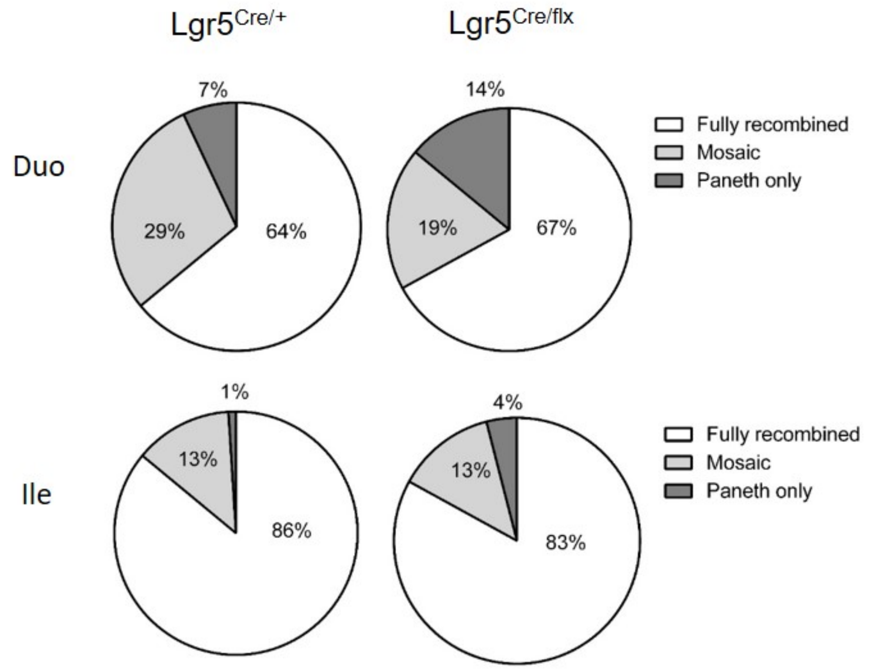

**Figure S4. Transcriptome analysis of Lgr5 ISC.**

A. ISC (eGFP[+ve]) cells from Lgr5-DTReGFP embryonic E18.5 and adult stages were sorted by FACS and RNA extracted was subject to RNAseq analysis. Heatmap of differentially regulated genes (fold change x1.5, FDR 0.1) in two independent samples (S1, S2) of E18.5 HE as compared to two independent samples (S1, S2) of adult ISCs from duodenum (Duo) and ileum (Ile). Selected genes are evidenced.
B. Graph showing relative expression of the Wnt ligands receptors/co-receptors (Fzd, Lrp, Gpc, Sdc) at different developmental stages E16.5, E18.5 and adult HEs based on the RNAseq data.

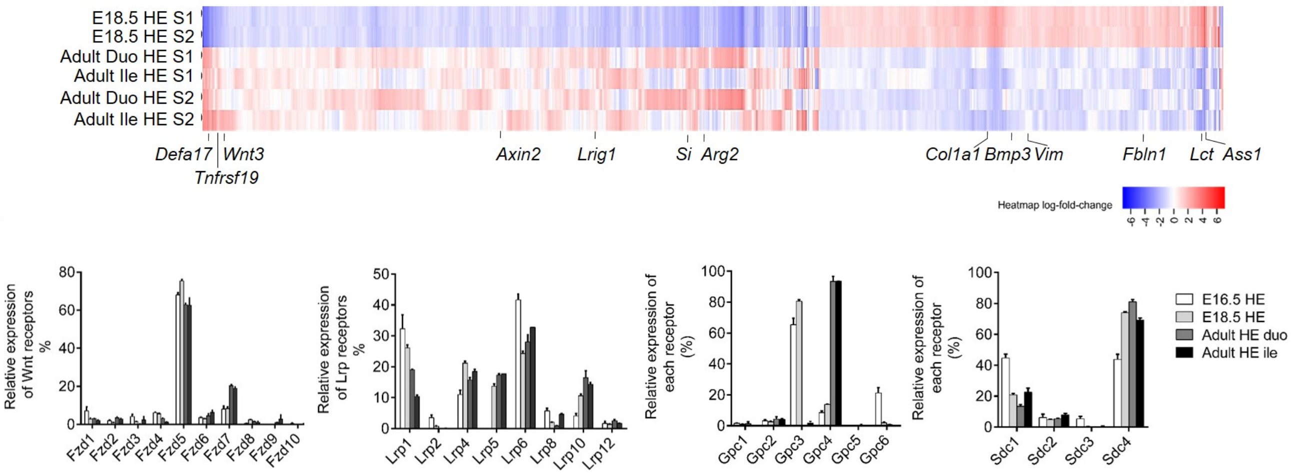

**Figure S5. Stemness is preserved in Lgr5-deficient organoids.**

A. Gene expression analysis by qRT-PCR of the indicated differentiation markers in Lgr5-DTReGFP WT and KO organoids cultured at the indicated Rspondin 1 concentrations after 5 days of culture. Each dot indicates the value for an organoid culture originating from a given embryo.
Data are represented as means ± sem. Analyzed by Kruskall-Wallis test followed by Dunns multiple comparison test.
B. Representative pictures showing Lgr5-DTReGFP WT and KO organoid cultures after 22 replatings. Scale bars, 50 µm.
C. Gene expression analysis by qRT-PCR of the indicated stem cell markers in Lgr5-DTReGFP WT and KO organoids after 5 days of culture at the indicated passages (pass 2 and pass 12: n= 7 WT, 9 KO; pass 22: n= 2 WT and 5 KO). Values are normalized to the WTs at passage 2.

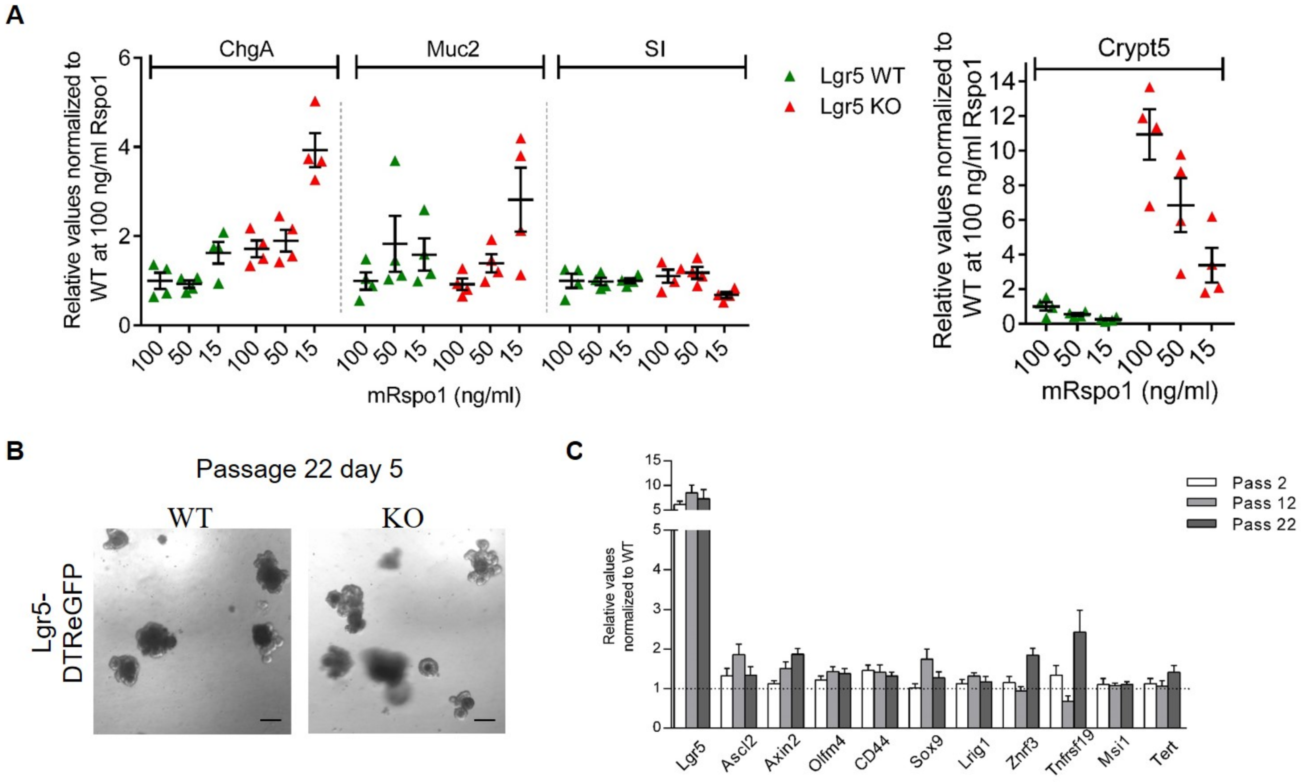
