## Supplementary material for "Lgr5 Controls Extracellular Matrix Production By Stem Cells In The Developing Intestine": Table S1 list of qPCR primers

| Gene | Forward primer | Reverse primer |
|---|---|---|
| Rpl13 | CCCGTGGCGATTGTGAA | TCATTGTCCTTCTGTGCAGGTT |
| gapdh | TTAGCCCCCCTGGCCAAGG | CTTACTCCTTGGAGGCCATG |
| axin2 | TGACTCTCCTTCCAGATCCCA | TGCCCACACTAGGCTGACA |
| chga | TCCCCACTGCAGCATCCAGTTC | CCTTCAGACGGCAGAGCTTCGG |
| lgr5 | CCTACTCGAAGACTTACCCAGT | GCATTGGGGTGAATGATAGCA |
| Tnfrsf19 | ACTGCATTTGGACGATCCTGG | CCAAAGAATGTGAACTGTAAACACC |
| Ascl2 | AAGCACACCTTGACTGGTACG | AAGTGGACGTTTGCACCTTCA |
| olfm4 | CAGCTGCCTGGTTGCCTCCG | GGCAGGTCCCATGGCTGTCC |
| Si | TTCAAGAAATCACAACATTCAATTTACTAG | CTAAAACTTTCTTTGACATTTGAGCAA |
| Muc2 | ATGCCCACCTCCTCAAAGAC | GTAGTTTCCGTTGGAACAGTGAA |
| Hopx | GTGCCTGCGATCTTGGTGGCT | GCCTGACCTTACGTCTGTCCCG |
| Tert | TGAGAGCAGCAGCAGCCTGT | TAGGCTGGAGCCCTGGGGGA |
| Znrf3 | GAACTTGACTGGCCTCCACCT | AAGAGGAAGCATGGTGAAAGCAC |
| Lyz1 | GAGACCGAAGCACCGACTATG | CGGTTTTGACATTGTGTTCGC |
| Mmp7 | ATGGCAGCTATGCAGCTCACCC | CCATATAACTTCTGAATGCCTGC |
| Crypt4 | AGAGACTAAAACTGAGGAGCAGC | CGGCGGGGGCAGCAGTA |
| crypt5 | AGAGACTAAAACTGAGGAGCAGC | GCAGCAGAATACGAAAGT |
| ywhaz | TGCAACGATCTACTGTCTCTTTTG | CGGTAGTAGTCACCCTTCATTTTCA |
